# Mitochondrial DNA Repair in an *Arabidopsis thaliana* Uracil N-Glycosylase Mutant

**DOI:** 10.1101/427500

**Authors:** Emily Wynn, Emma Purfeerst, Alan Christensen

## Abstract

Substitution rates in plant mitochondrial genes are extremely low, indicating strong selective pressure as well as efficient repair. Plant mitochondria possess base excision repair pathways, however, many repair pathways such as nucleotide excision repair and mismatch repair appear to be absent. In the absence of these pathways, many DNA lesions must be repaired by a different mechanism. To test the hypothesis that double-strand break repair (DSBR) is that mechanism, we maintained independent self-crossing lineages of plants deficient in uracil-N-glycosylase (UNG) for 11 generations to determine the repair outcomes when that pathway is missing. Surprisingly, no single nucleotide polymorphisms (SNPs) were fixed in any line in generation 11. The pattern of heteroplasmic SNPs was also unaltered through 11 generations. When the rate of cytosine deamination was increased by mitochondrial expression of the cytosine deaminase APOBEC3G, there was an increase in heteroplasmic SNPs, but only in mature leaves. Clearly DNA maintenance in reproductive meristem mitochondria is very effective in the absence of UNG, while mitochondrial genomes in differentiated tissue are maintained through a different mechanism, or not at all. Several genes involved in DSBR are upregulated in the absence of UNG, indicating that double strand break repair is a general system of repair in plant mitochondria. It is important to note that the developmental stage of tissues is critically important for these types of experiments.

## 1. Introduction

Plant mitochondrial genomes have very low base substitution rates, while also expanding and rearranging rapidly [1-5]. The low substitution rate and the high rearrangement rate of plant mitochondria can be explained by selection and the specific DNA damage repair mechanisms available. These mechanisms can also account for the genome expansions often found in land plant mitochondria [6]. The low nonsynonymous substitution rates in protein coding genes indicate that selective pressure to maintain the genes is high, and the low synonymous substitution rates indicate that the DNA repair mechanisms are very accurate [7, 8]. Despite the low mutation rate of mitochondrial genes over evolutionary time, mitochondrial genomes in mature cells accumulate DNA damage that is not repaired [9]. This indicates that there are fundamental differences between DNA maintenance in genomes meant to be passed on to the next generation and genomes that are not. In meristematic cells, mitochondria fuse together to form a large mitochondrion [10]. This fusion brings mitochondrial genomes together for genome replication, but also ensures that there is a homologous template available for DNA repair. These meristematic cells eventually produce the reproductive tissue of a plant; from embryogenesis to egg cell production the mitochondrial genomes inherited from parents and passed down to offspring will have homologous templates available to them [11].

Much less is known about the multiple pathways of DNA repair in plant mitochondria than in other systems, such as the nucleus. So far, there is no evidence of nucleotide excision repair (NER), nor mismatch repair (MMR) in plant mitochondria [12, 13]. It has been hypothesized that in plant mitochondria, the types of DNA damage that are usually repaired through NER and MMR are repaired through double-strand break repair (DSBR) [14, 15]. Plant mitochondria do have the nuclear-encoded base excision repair (BER) pathway enzyme Uracil DNA glycosylase (UNG) [12]. UNG is an enzyme that can recognize and bind to uracil in DNA and begin the process of base excision repair by enzymatically excising uracil (U) from single stranded or double stranded DNA [16]. Uracil can appear in a DNA strand due to the spontaneous deamination of cytosine, or by the misincorporation of dUTP during replication [17]. Unrepaired uracil in DNA can lead to G-C to A-T transitions within the genome.

In light of the apparent absences of NER and MMR in plant mitochondria, it is possible that many lesions, including mismatches, are repaired by creating double-strand breaks and using a template to repair both strands. Our hypothesis is that DSBR accounts for most of the repair in meristematic plant mitochondria, and both error-prone and accurate subtypes of DSBR lead to the observed patterns of genome evolution [18]. One way of testing this is to eliminate the pathway of uracil base excision repair and ask if the G-U mispairs that occur by spontaneous deamination are repaired, and if so, are instead repaired by DSBR. In this work we examine an *Arabidopsis thaliana ung* knockout line and investigate the effects on the mitochondrial genome over many generations. To disrupt the genome further, we express the cytidine deaminase APOBEC3G in the *Arabidopsis* mitochondria (MTP-A3G) to increase the rate of cytosine deamination and accelerate DNA damage.

One of the hallmarks of DSBR in plant mitochondria is the effect on the non-tandem repeats that exist in virtually all plant mitochondria [19]. The *Arabidopsis thaliana* mitochondrial genome contains two pairs of very large repeats (4.2 and 6.6kb) that commonly undergo recombination [20-22] producing multiple isoforms of the genome. The mitochondrial genome also contains many non-tandem repeats between 50 and 1000 base pairs,[19, 22-24]. In wild type plants, these repeats recombine at very low rates, but they have been shown to recombine with ectopic repeat copies at higher rates in several mutants in DSBR-related genes, such as *msh1* and *reca3* [25-27]. Thus, genome dynamics around non-tandem repeats can be an indicator of increased DSBs. In this work we show that a loss of uracil base excision repair leads to alterations in repeat dynamics.

Numerous proteins known to be involved in the processing of plant mitochondrial DSBs have been characterized. Plants lacking the activity of mitochondrially targeted *recA* homologs have been shown to be deficient in DSBR [26, 28]. In addition, it has been hypothesized that the plant MSH1 protein may be involved in binding to DNA lesions and initiating DSBs [14, 15]. The MSH1 protein contains a mismatch binding domain fused to a GIY-YIG type endonuclease domain which may be able to make DSBs [29, 30]. In this work we provide evidence that in the absence of mitochondrial UNG activity, several genes involved in DSBR, including *MSH1*, are transcriptionally upregulated, providing a possible explanation for the increased DSBR. We also provide additional evidence to support the hypothesis that mitochondrial DNA maintenance is abandoned in non-meristematic tissue [31], calling attention to the need to closely control for age and developmental state in experiments involving the mitochondrial genome.

## 2. Results

### 2.1. Lack of UNG activity in mutants

It has previously been reported that cell extracts of the *Arabidopsis thaliana ung* T-DNA insertion strain used in this experiment, GK-440E07 (ABRC seed stock CS308282), show no uracil glycosylase activity [12]. To increase the rate of cytosine deamination in the mitochondrial genome and show that effects of the UNG knockout on mitochondrial mutation rates could be detected, the catalytic domain of the human APOBEC3G–CTD 2K3A cytidine deaminase (A3G) [32] was expressed under the control of the Ubiquitin-10 promoter [33] in both wild-type and *ung Arabidopsis thaliana* lines and targeted to the mitochondria by an amino-terminal fusion of the 62 amino acid mitochondrial targeting peptide (MTP) from the Alternative Oxidase 1A protein. Fluorescence microscopy of *Arabidopsis thaliana* expressing an MTP-A3G-GFP fusion shows that the MTP-A3G construct is expressed and targeted to the mitochondria (Supplemental Figure S1).

We expected that in the absence of UNG there would be an increase in G-C to A-T substitution mutations. To test this prediction, we sequenced a wild-type Arabidopsis plant (Col-0), a wild-type Arabidopsis plant expressing the MTP-A3G construct (Col-0 MTP-A3G) and a *ung* plant expressing the MTP-A3G construct (*ung* MTP-A3G) using an Illumina Hi-Seq4000 system. Mitochondrial sequence reads from these plants were aligned to the Columbia-0 reference genome (modified as described in Materials and Methods) using BWA-MEM [34] and single nucleotide polymorphisms were identified using VarDict [35].

There were no SNPs that reached fixation (an allele frequency of 1) in any plant. Mitochondrial genomes are not diploid; each cell can have many copies of the mitochondrial genome. Therefore, it is possible that an individual plant could accumulate low frequency mutations in some of the mitochondrial genomes in the cell. VarDict was used to detect heteroplasmic SNPs at allele frequencies as low as 0.01. VarDict’s sensitivity in calling low frequency SNPs scales with depth of coverage and quality of the sample, so it is not possible to directly compare heteroplasmic mutation rates in samples with different depths of coverage. However, because the activity of the UNG protein is specific to uracil, the absence of the UNG protein should not have any effect on mutation rates other than G-C to A-T transitions. Comparing the numbers of G-C to A-T transitions to all other substitutions should reveal if the rate of mutations that can be repaired by UNG is elevated compared to the background rate. If the *ung* MTP-A3G line is accumulating G-C to A-T transitions at a faster rate than the Col-0 MTP-A3G line, we would expect to see that as an increased ratio of G-C to A-T transitions compared to other mutation types. Complicating the analysis, significant portions of the *A. thaliana* mitochondrial genome have been duplicated in the nucleus, forming regions called NuMTs. Mutations in the NuMTs might appear to be low frequency SNPs in the mitochondrial genome, confounding the results. However, these mutations are likely to be shared in the common nuclear background of all our lines. To avoid attributing SNPs in NuMTs to the mitochondrial genome, only those SNPs unique to individual plant lines were used in this comparison. The Col-0 plant had a heteroplasmic GC-AT/total SNPs ratio of 0.073, the Col-0 MTP-A3G plant had a heteroplasmic GC-AT/total SNPs ratio of 0.44, while the *ung* MTP-A3G plant had a heteroplasmic GC-AT/total SNPs ratio of 0.61 (Table 1). Therefore, when the rate of cytosine deamination is increased by the activity of APOBEC3G, Arabidopsis plants accumulate GC-AT SNPs, and *ung* plants accumulate GC-AT SNPs faster than wild-type plants. Our computational pipeline is able to detect both MTP-A3G activity and the effect of the *ung* knockout.

**Table 1.** Heteroplasmic mitochondrial SNPs in Col-0 wild-type, generation 10 *ung* mutant lines, Col-0 MTP-A3G, and *ung* MTP-A3G. SNPs were called using VarDict as described in Methods. SNP counts are shown for the entire mitochondrial genome. For the full spectrum of SNP types, including allele frequencies, see supplemental file 2.

| Sample | GC-AT<br>SNPs | Total<br>SNPs | GC-AT<br>/Total<br>SNP Ratio |
| --- | --- | --- | --- |
| Col-0 | 17 | 234 | 0.073 |
| *ung*10 115 | 9 | 96 | 0.094 |
| *ung*10 159 | 12 | 146 | 0.082 |
| *ung*10 163 | 69 | 1261 | 0.055 |
| *ung*10 176 | 42 | 615 | 0.068 |
| *ung*10 198 | 9 | 73 | 0.12 |
| *ung*10 201 | 6 | 126 | 0.048 |
| *ung*10 203 | 8 | 101 | 0.079 |
| Col-0 MTP-A3G | 78 | 177 | 0.44 |
| *ung* MTP-A3G | 76 | 124 | 0.61 |

### 2.2. Mutation accumulation in the absence of UNG

To determine the effects of the *ung* knockout across multiple generations, we performed a mutation accumulation study [36]. We chose 23 different *ung* homozygous plants derived from one hemizygous parent. These 23 plants were designated as generation 1 *ung* and allowed to self-cross. The next generation was derived by single-seed descent from each line, and this was repeated until generation 10 *ung* plants were obtained. Leaf tissue and progeny seeds from each line were kept at each generation.

The leaf tissue from generation 10 of the *ung* mutation accumulation lines and a wild-type Col-0 were sequenced and analyzed with VarDict as described above. Similar to the MTP-A3G plants, there were no SNPs in any of our *ung* mutation accumulation lines that had reached fixation (an allele frequency of one). In contrast, there was no relative increase in the ratios of GC-AT/total SNPs between the *ung* lines and Col-0 (see Table 1). Because detection of low frequency SNPs depends on read depth, we only report the 7 *ung* samples with an average mitochondrial read depth above 125x for this comparison. In the absence of a functional UNG protein and under normal greenhouse physiological conditions, plant mitochondria do not accumulate cytosine deamination mutations at an increased rate.

### 2.3 Nuclear Mutation Accumulation

UNG is the only Uracil-N-Glycosylase in *Arabidopsis thaliana* and may be active in the nucleus as well as the mitochondria [12]. To test for nuclear mutations due to the absence of UNG, sequences were aligned to the Columbia-0 reference genome using BWA-MEM and single nucleotide polymorphisms were identified using Bcftools Call [37]. There is more variability in nuclear mutation ratios than mitochondrial due to the low total number of SNPs detected, however the *ung* mutation accumulation lines do not have an elevated G-C to A-T mutation rate compared to wild-type(Table 2).

**Table 2:** Nuclear SNPs in Col-0 wild-type, *ung* mutant lines, Col-0 MTP-A3G, and *ung* MTP-A3G. SNPs were called using Bcftools Call as described in Methods. SNP counts are shown for 5Mb regions of each chromosome, excluding chromosome 2. For individual data on each chromosome see supplemental file 2.

| Sample | GC-AT | Total | GC-AT |
| --- | --- | --- | --- |

|  | SNPs | SNPs | /Total<br>SNP Ratio |
| --- | --- | --- | --- |
| Col-0 | 1 | 3 | 0.33 |
| *ung10 115* | 1 | 16 | 0.063 |
| *ung10 159* | 0 | 8 | 0 |
| *ung10 163* | 5 | 10 | 0.5 |
| *ung10 176* | 0 | 5 | 0 |
| *ung10 198* | 3 | 14 | 0.21 |
| *ung10 201* | 4 | 21 | 0.19 |
| *ung10 203* | 3 | 11 | 0.27 |
| Col-0 MTP-A3G | 1 | 3 | 0.33 |
| *ung* MTP-A3G | 0 | 10 | 0 |

### 2.4 Alternative Repair Pathway Genes

Because the *ung* mutants show increased double-strand break repair but not an increase of G-C to A-T transition mutations, we infer that the inevitable appearance of uracil in the DNA is repaired via conversion of a G-U pair to a double-strand break and efficiently repaired by the DSBR pathway. If this is true, genes involved in the DSBR processes of breakage, homology surveillance and strand invasion in mitochondria will be up-regulated in *ung* mutants. To test this hypothesis, we assayed transcript levels of several candidate genes known to be involved in DSBR [13, 23, 25-28, 38-41] in *ung* lines compared to wild-type using RT-PCR. *MSH1* and *RECA2* were significantly upregulated in *ung* lines (*MSH1*: 5.60-fold increase, unpaired T-test p<0.05. *RECA2*: 3.19-fold increase, unpaired T-test p<0.05 – see Figure 2). The single-strand binding protein gene *OSB1* was also measurably upregulated in *ung* lines (3.07-fold increase, unpaired T-test p=0.053). *RECA3, SSB*, and *WHY2* showed no differential expression compared to wild-type (unpaired T-test p>0.05).

**Figure 1:**
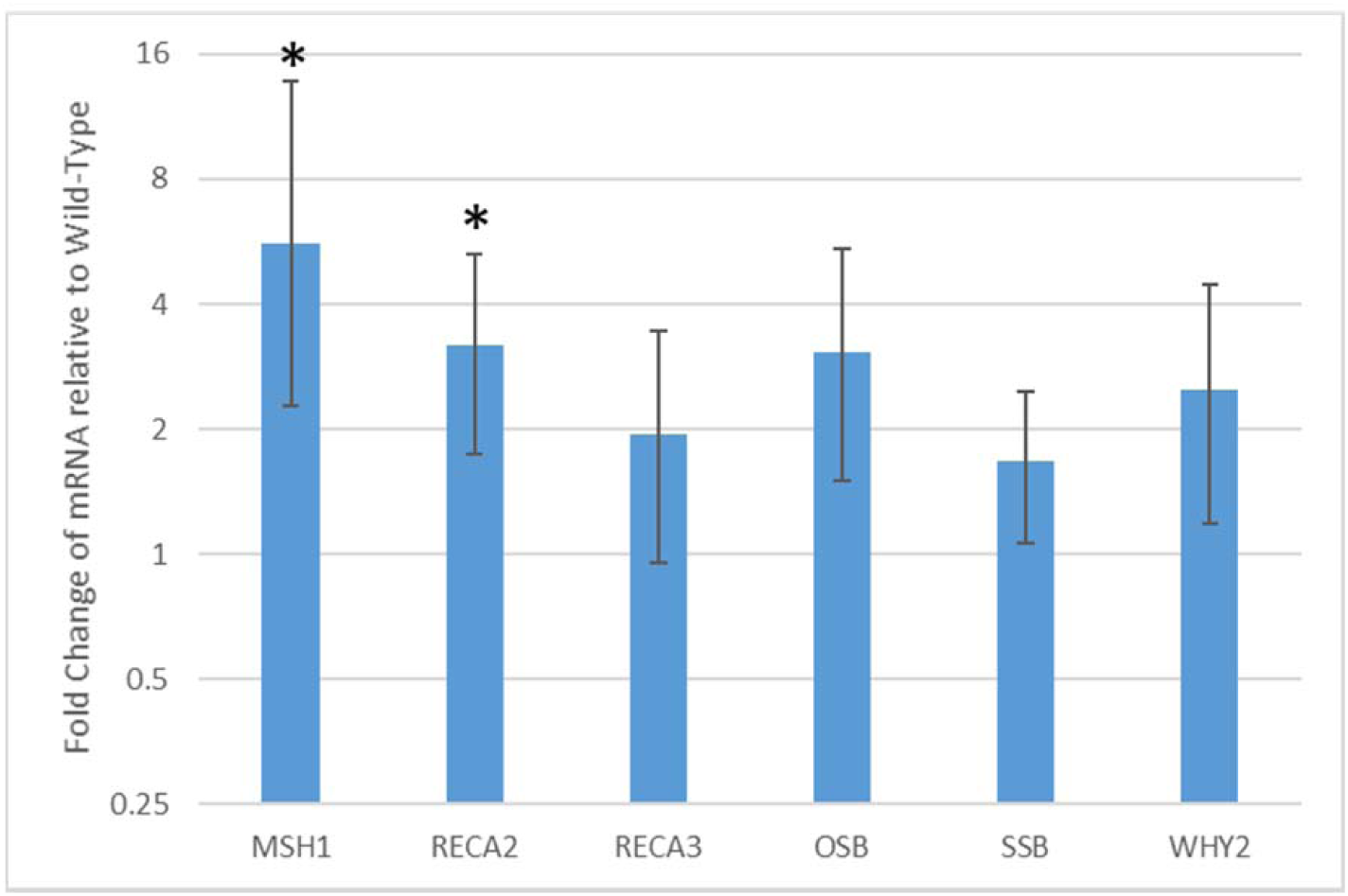
Quantitative RT-PCR assays of enzymes involved in DSBR in *ung* lines relative to wild-type. Fold change in transcript level is shown on the Y-axis. Error bars are standard deviation of three biological replicates. *MSH1* and *RECA2* are significantly transcriptionally upregulated in *ung* lines relative to wild-type (5.60-fold increase and 3.19-fold increase, respectively. Unpaired, 2-tailed Student’s t-test, * indicates p<0.05). *OSB1* is nearly significantly upregulated in *ung* lines relative to wild-type (3.07-fold increase. Unpaired T-test p=0.053).

**Figure 2:**
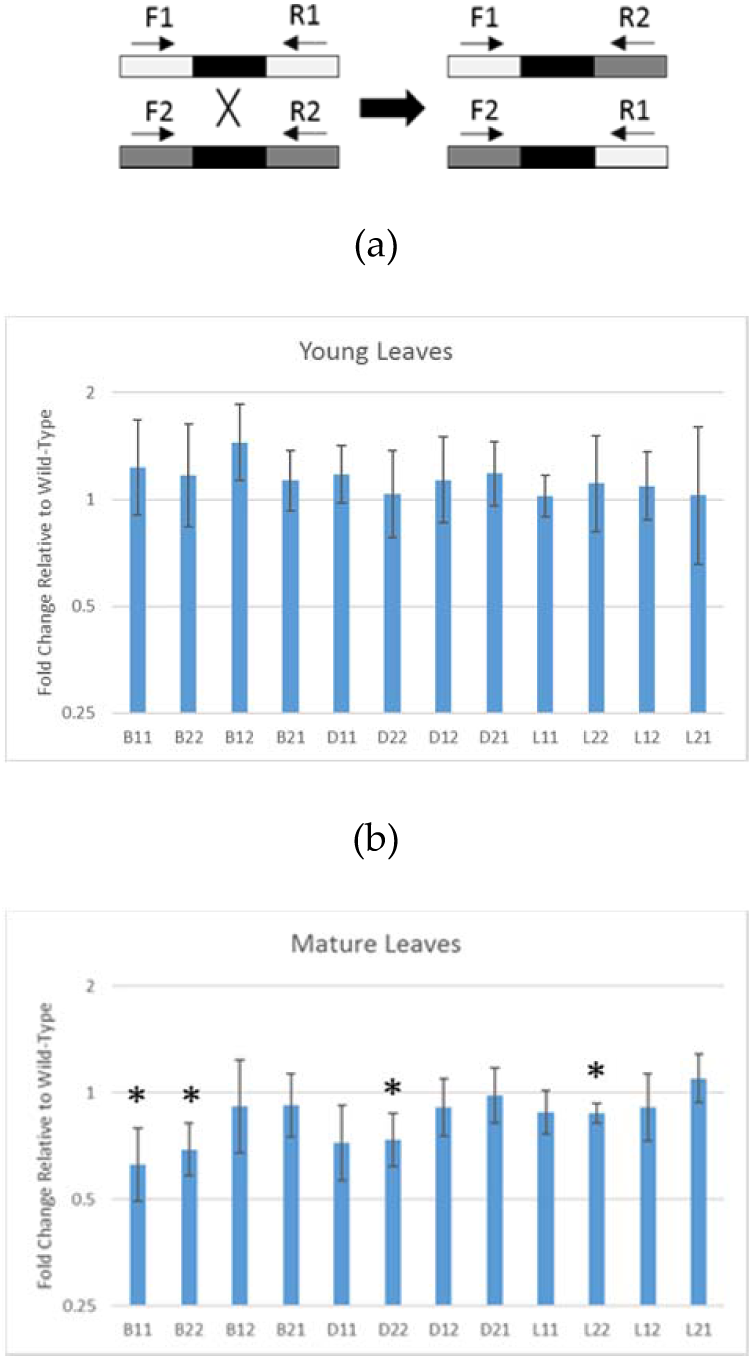
qPCR analysis of intermediate repeat recombination in *ung* lines compared to wild-type. Recombination at intermediate repeats is an indicator of increased double strand breaks in plant mitochondrial genomes. **(a)** Primer scheme for detecting parental and recombinant repeats. Using different combinations of primers that anneal to the unique sequence flanking the repeats, either parental type (1/1 and 2/2) or recombinant type (1/2 and 2/1) repeats can be amplified **(b)** Fold change of intermediate repeats in young leaves of *ung* lines relative to wild-type. Error bars are standard deviation of three biological replicates. **(c)** Fold change of intermediate repeats in mature leaves of *ung* lines relative to wild-type. Error bars are standard deviation of three biological replicates. B1/1, B2/2, D2/2, and L2/2 show significant reduction in copy number (unpaired, 2-tailed Student’s t-test, * indicates p<0.05)

### 2.5 Increased Double-strand break repair

If most DNA damage in plant mitochondria is repaired by double-strand break repair (DSBR), supplemented by base excision repair [12], then in the absence of the Uracil-N-glycosylase (UNG) pathway we predict an increase in DSBR. To find evidence of this we used quantitative PCR (qPCR) to assay crossing over between identical non-tandem repeats because changes in the dynamics around these repeats is indicative of changes in DNA processing at double-strand breaks [26, 27, 41]. Different combinations of primers in the unique sequences flanking the repeats allow us to determine the relative copy numbers of parental-type repeats and low frequency recombinants (Figure 2a). The mitochondrial genes *cox2* and *rrn18* were used to standardize relative amplification between lines. We and others [24, 41] have found that some of the non-tandem repeats are well-suited for qPCR analysis and are sensitive indicators of ectopic recombination, increasing in repair-defective mutants. We analyzed the three repeats known as Repeats B, D, and L [23] in both young leaves and mature leaves. In young leaves, there is no significant difference in the amounts of parental or recombinant forms between *ung* lines and Col-0 (Figure 2b). In mature leaves, all three repeats show significant reductions in the parental 2/2 form, while repeat B also shows a reduction in the parental 1/1 form (unpaired T-test p<0.05, Figure 2c). There is a difference in genome dynamics around non-tandem repeats in young leaves compared to old leaves, indicating a difference in the way these genomes are maintained.

### 2.6 Transmission of SNPs across generations

To determine if any heteroplasmic SNPs are passed on to the next generation, two progeny of each of the wild-type, *ung*, MTP-A3G, and *ung* MTP-A3G plants that were sequenced above were planted. Leaves were collected from each plant when it was 17 days old (young leaf) and again when it was 36 days old (mature leaf). Both the young and mature leaves of each plant were sequenced and analyzed as described above. Only 2 heteroplasmic SNPs could be traced from a parent plant to both progeny, and 42 heteroplasmic SNPs could be traced from a parent plant to one progeny (Supplemental File 2). Interestingly, 115 heteroplasmic SNPs were detected in both offspring but not the parent plant. It is possible that heteroplasmic mutations that occur in reproductive tissue after the parental tissue had been collected could be passed on to the progeny. However, only 17 of these heteroplasmic SNPs are found in the mature tissue of both progeny, indicating that even if a heteroplasmic SNP is passed on to a future generation, it is likely to be removed from the mitochondrial population before reproduction by genetic drift or gene conversion. In fact, of the 13,914 heteroplasmic SNPs that were detected in young tissue across all samples, only 31 were detected in the mature tissue of the same plant. The overwhelming majority of heteroplasmic SNPs arose in mitochondria in non-meristematic differentiated tissue.

### 2.7 SNP accumulation in young vs mature leaves

To confirm that the effects of the UNG knockout and the expression of APOBEC3G are consistent, the progeny of the wild-type, *ung*, MTP-A3G, and *ung* MTP-A3G plants were analyzed and the ratio of heteroplasmic GC-AT to total heteroplasmic SNPs was compared as described above. In mature leaves, the results were similar to the previous generation: both the *ung* MTP-A3G and Col-0 MTP-A3G samples had increased GC-AT SNPs compared to the *ung* and Col-0. Interestingly, in young leaves, neither the *ung* MTP-A3G nor the Col-0 MTP-A3G samples had increased GC-AT SNPs (See Table 3). This indicates that the processes of mitochondrial genome maintenance is more efficient at repairing DNA damage in young leaves.

**Table 3:**
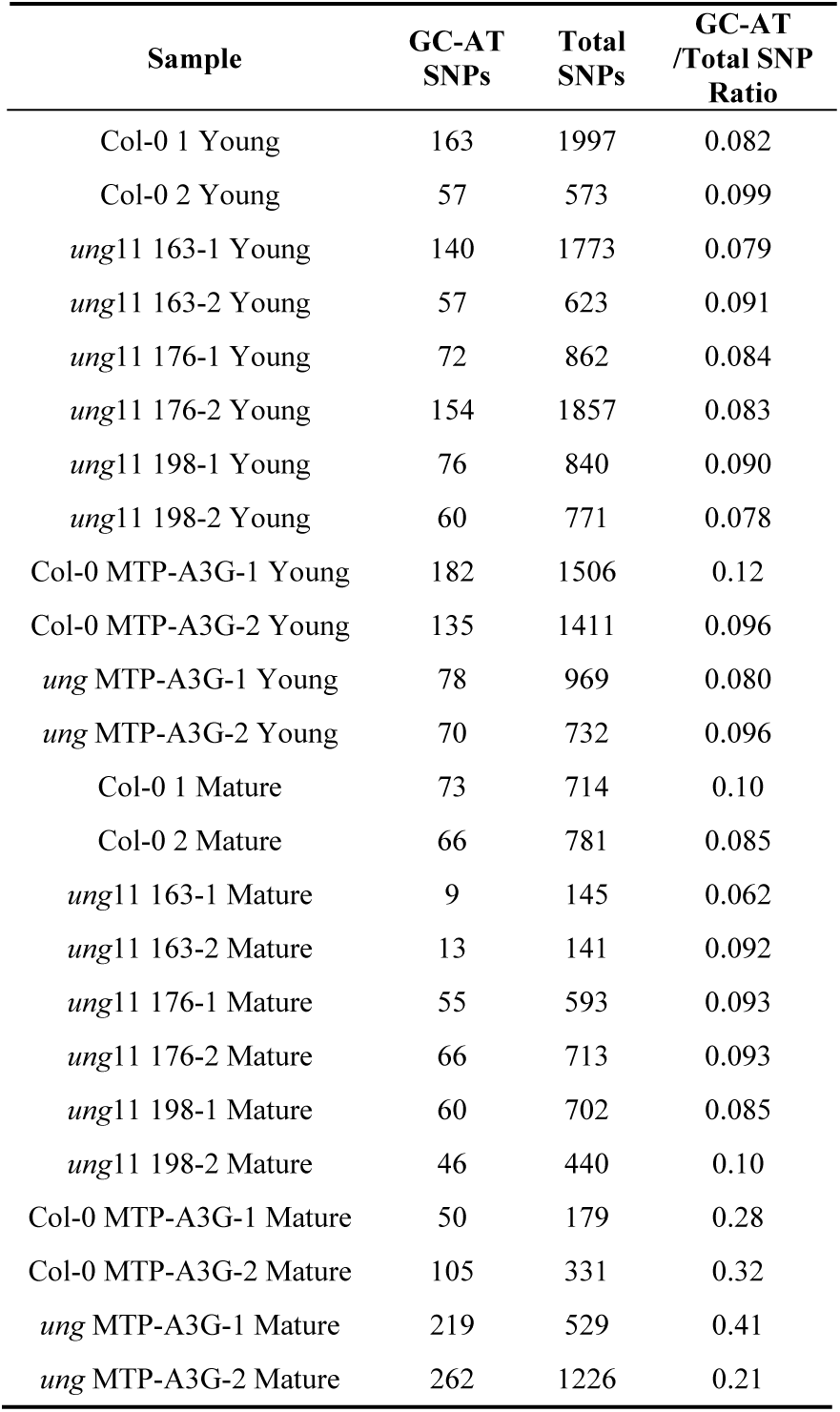
Heteroplasmic mitochondrial SNPs in the next generation of Col-0 wild-type, *ung mutant lines, Col-0 MTP-A3G, and ung MTP-A3G*. SNPs were called using VarDict as described in Methods. SNP counts are shown for the entire mitochondrial genome. For the full spectrum of SNP types, including allele frequencies, see Supplemental file 2.

### 2.8 Quality Control of DNA Library Preparation

A common source of error when calling low frequency SNPs is oxidative damage during library preparation [42]. This oxidative damage affects Guanines, and the effects of this damage can be measured by comparing the ratio of G to T mutations between the R1 paired-end read and the R2 paired-end read. A Global Imbalance Value for G to T changes (GIV_G_T_) above 1.5 indicates DNA damage during library preparation, while a GIV_G_T_ below 1.5 indicates little damage during library preparation. All of the *ung* generation 10 libraries had a GIV_G_T_ above 1.5. However, all other samples, including all *ung* generation 11 libraries, had a GIV_G_T_ below 1.5 (see Supplemental file 2). For all data reported in this study, there is little difference between *ung* generation 10 and *ung* generation 11 samples. Any damage caused during library preparation of *ung* generation 10 samples does not change the outcome or interpretation of this study.

## 3. Discussion

In mitochondria as well as in the nucleus and chloroplast, cytosine is subject to deamination to uracil. This could potentially lead to transition mutations and is dealt with by a specialized base excision repair pathway. The first step in this pathway is hydrolysis of the glycosidic bond by the enzyme Uracil-N-glycosylase (UNG), leaving behind an abasic site [16]. An AP endonuclease can then cut the DNA backbone, producing a 3’ OH and a 5’ dRP. Both DNA polymerases found in *A. thaliana* mitochondria, POL1A and POL1B, exhibit 5’-dRP lyase activity, allowing them to remove the 5’ dRP and polymerize a new nucleotide replacing the uracil [43]. In the absence of functional UNG protein, cytosine will still be deaminated in plant mitochondrial genomes, so efficient removal of uracil must be through a different repair mechanism, most likely DSBR [14, 15]. We have found that in *ung* mutant lines, there is an increase in the expression of genes known to be involved in DSBR, and significant changes in the relative abundance of parental and recombinant forms of intermediate repeats, consistent with this hypothesis.

We have shown that when cytosine deamination is increased by the expression of the APOBEC3G cytidine deaminase in plant mitochondria, *ung* lines accumulate more G-C to A-T transitions in mature leaves, than does wild-type. Surprisingly, we have also found that under normal cellular conditions, without the added deamination activity of APOBEC3G, *ung* lines do not accumulate G-C to A-T transition mutations at a higher rate than wild-type. This finding is particularly surprising given the presumed bottlenecking of mitochondrial genomes during female gametogenesis and given the deliberate bottleneck in the experimental design of single-seed descent for 11 generations. This finding supports the hypothesis that plant mitochondria have a very efficient alternative damage surveillance system that can prevent G-C to A-T transitions from becoming fixed in the meristematic mitochondrial population.

The angiosperm MSH1 protein consists of a DNA mismatch binding domain fused to a double-stranded DNA endonuclease domain [1, 21] Although mainly characterized for its role in recombination surveillance [35], MSH1 is a good candidate for a protein that may be able to recognize and bind to various DNA lesions and make DSBs near the site of the lesion, thus funneling these types of damage into the DSBR pathway. With many mitochondria and many mitochondrial genomes in each cell there are numerous available templates for accurate repair of DSBs through homologous recombination, making this a plausible mechanism of genome maintenance. Here we show that in *ung* lines, *MSH1* is transcriptionally upregulated more than 5-fold compared to wild-type. This is consistent with the hypothesis that MSH1 initiates repair in plant mitochondria by creating a double-strand break at G-U pairs, and possibly other mismatches and damaged bases.

Several other proteins involved in processing plant mitochondrial DSBs have been characterized. The RECA homologs, RECA2 and RECA3, are homology search and strand invasion proteins [26-28, 40, 44-46]. The two mitochondrial RECAs share much sequence similarity, however RECA2 is dual targeted to both the mitochondria and the plastids, while RECA3 is found only in the mitochondria [26, 27]. RECA3 also lacks a C-terminal motif present on RECA2 and most other homologs. This motif has been shown to modulate the ability of RECA proteins to displace competing ssDNA binding proteins in *E. coli* [47]. Arabidopsis *reca2* mutants are seedling lethal and both *reca2* and *reca3* lines show increased ectopic recombination at intermediate repeats [26]. Arabidopsis RECA2 has functional properties that RECA3 cannot perform, such as complementing a bacterial *recA* mutant during the repair of UV-C induced DNA lesions [20]. Here we show that in *ung* lines, *RECA2* is transcriptionally upregulated more than 3-fold compared to the wild-type. However, *RECA3* is not upregulated in *ung* lines. Responding to MSH1-initiated DSBs may be one of the functions unique to RECA2. The increased expression of *RECA2* in the absence of a functional UNG protein is further evidence that uracil arising in DNA may be repaired through the mitochondrial DSBR pathway.

The ssDNA binding protein OSB1’s transcript is upregulated over 3-fold. At a double strand break, OSB1 competitively binds to ssDNA and recruits the RECA proteins to promote the repair of a double strand break by a homologous template and avoid the error-prone microhomology-mediated end-joining pathway [48].

We also tested the differential expression of other genes known to be involved in processing mitochondrial DSBs. The single stranded binding protein genes *WHY2* and *SSB* were not found to be differentially expressed at the transcript level compared to wild-type. The presence of different ssDNA binding proteins influences which pathway of DSBR a break is repaired by [48]. Increased amounts of WHY2 and SSB may not be needed for accurate repair of induced DSBs in the *ung* lines.

The specific patterns of recombination at mitochondrial intermediate repeats are different between wild-type, *ung* mutants, and DSBR mutants. In *msh1* lines, there is an increase in repeat recombination likely due to relaxed homology surveillance in the absence of the MSH1 protein [27]. In mutant lines of ssDNA binding proteins involved in DSBR, such as *recA2, recA3*, and *osb1* [26, 49], there is an increase in repeat recombination due to differences in the way DNA ends are handled in the absence of these ssDNA binding proteins. In *ung* lines, the mitochondrial recombination machinery is still intact, so any differences in repeat recombination between *ung* lines and wild-type are not due to differences in processing the DSB, but due to the increase in the amount of DSBs in the absence of UNG.

In young leaves, there is no significant difference in recombination at intermediate repeats between *ung* lines and wild-type, while in mature leaves, *ung* lines show a reduction in parental type repeats compared to wild-type. This could indicate that there is an increase in double strand breaks and an increase in attempted DSBR by break-induced replication at intermediate repeats or be an indication of degradation of mtDNA as differentiated tissue ages.

Plant mitochondrial genomes likely replicate by recombination-dependent replication (RDR) [50]. Most organellar genome replication occurs in meristematic tissue, where mitochondria fuse together to form a large, reticulate mitochondrion [10] This mitochondrial fusion provides a means to homogenize mtDNA by gene conversion, and repair lesions through homologous recombination [51]. Accurate repair of a uracil by homologous recombination would not be expected to change repeat dynamics. As cells differentiate and age, organellar genomes degrade [31]. Organellar genomes in non-reproductive tissue can be “abandoned” rather than repaired, reducing the metabolic cost of DNA repair [31]. In a mature cell, an attempt to repair uracil in the mitochondrial genome could lead to degradation of the DNA and changes in repeat dynamics if a double-strand break is initiated without a homologous template available. Clearly there is a difference in mitochondrial DNA maintenance in mature cells compared to young cells, either due to a lack of DNA repair in mature mitochondria, or a difference in DNA repair mechanism.

To determine the outcomes of genomic uracil in the absence of a functional UNG protein, we sequenced the genomes of several *ung* lines. No fixed mutations of any kind were found in *ung* lines, even after 11 generations of self-crossing. Low frequency heteroplasmic SNPs were found in both wild-type and *ung* lines, but *ung* lines showed no difference in the ratio of G-C to A-T transitions to other mutation types when compared to wild-type. When the rate of cytosine deamination was increased with the expression of the APOBEC3G deaminase, there was an increase in G-C to A-T transitions, but only in mature leaves. This is consistent with the idea of abandonment and is evidence that in mitochondrial genomes that have not been abandoned, there is an efficient and accurate system of non-specific repair.

Clearly the double-strand break repair pathway in plant mitochondria can repair uracil in DNA sufficiently to prevent mutation accumulation in the absence of the UNG protein. Why then has the BER pathway been conserved in plant mitochondria while NER and MMR have apparently been lost? DSBR protects the genome efficiently from mutations being inherited by the next generation (see Table 3). There may still be selection to maintain mitochondrial BER to reduce the rate of mitochondrial genome abandonment and degradation in aging tissues. Throughout the evolutionary history of *Arabidopsis thaliana* and into the present, wild growing plants are exposed to a range of growth conditions and stresses that experimental plants in a greenhouse avoid. The rate of spontaneous cytosine deamination increases with increasing temperature [52, 53], so DSBR alone may not be able repair the extent of uracil found in DNA across the range of temperatures a wild plant would experience, providing the selective pressure to maintain a distinct BER pathway in plant mitochondria. If DSBR activity is reduced or lost as leaf tissue ages, there may also be a selective advantage to the plant of maintaining BER in mature leaves so they can continue to perform intermediary metabolism even as they age.

Here we have provided evidence that in the absence of a dedicated BER pathway, plants growing in greenhouse growth chamber conditions do not accumulate mitochondrial SNPs at an increased rate. Instead, DNA damage is accurately repaired by double-strand break repair which also causes an increase in ectopic recombination at identical non-tandem repeats. It has recently been shown that mice lacking a different mitochondrial BER protein, oxoguanine glycosylase, also do not accumulate mitochondrial SNPs [54]. Here we show that in plants base-excision repair by UNG is similarly unnecessary to prevent mitochondrial mutations in growth chamber conditions. Clearly DSBR is efficient and accurate, and the presence of the UNG pathway reduces ectopic recombination slightly and can successfully repair uracil in DNA even if the rate of cytosine deamination is increased. We have also found that in mature leaves, uracil mutations do occur, further confirming the hypothesis that organellar genomes are abandoned in terminally differentiated tissues [31] and emphasizing the need for considering the tissue age and type when interpreting experimental results on DNA replication, repair and recombination. Double strand break repair and recombination are important mechanisms in the evolution of plant mitochondrial genomes, but many key enzymes and steps in the repair pathway are still unknown. Further identification and characterization of these missing steps is sure to provide additional insight into the unique evolutionary dynamics of plant mitochondrial genomes.

## 4. Materials and Methods

### 4.1 Plant growth conditions

*Arabidopsis thaliana* Columbia-0 (Col-0) seeds were obtained from Lehle Seeds (Round Rock, TX, USA). *UNG* (AT3G18630) T-DNA insertion hemizygous lines were obtained from the Arabidopsis Biological Resource Center, line number CS308282. Hemizygous T-DNA lines were self-crossed to obtain homozygous lines (Genotyping primers: wild-type 5’-TGTCAAAGTCCTGCAATTCTTCTCACA-3’ and 5’-TCGTGCCATATCTTGCAGACCACA-3’, *ung* 5’-ATAATAACGCTGCGGACATCTACATTTT-3’ and 5’-ACTTGGAGAAGGTAAAGCAATTCA-3’). All plants were grown in walk-in growth chambers under a 16:8 light:dark schedule at 22°C. Plants grown on agar were surface sterilized and grown on 1x Murashige and Skoog Basal Medium (MSA) with Gamborg’s vitamins (Sigma) with 5μg/mL Nystatin Dihydrate to prevent fungal contamination.

### 4.2 Vector construction

The APOBEC3G gene [55]was synthesized by Life Technologies Gene Strings using *Arabidopsis thaliana* preferred codons and including the 62 amino acid mitochondrial targeting peptide (MTP) from Alternative Oxidase on the N-terminus of the translated protein. The MTP-A3G construct was cloned into the vector pUB-DEST (NCBI:taxid1298537) driven by the ubiquitin (UBQ10) promoter and transformed into wild-type and *ung Arabidopsis thaliana* plants by the *Agrobacterium* floral dip method [56]. To ensure proper mitochondrial targeting of the MTP-A3G construct, the construct was cloned into pK7FWG2 with a C-terminal GFP fusion [57]. *Arabidopsis thaliana* plants were again transformed by the *Agrobacterium* floral dip method and mitochondrial fluorescence was confirmed with confocal fluorescence microscopy.

### 4.3 RT-PCR

RNA was extracted from young leaves of plants grown on MSA during *ung* generation ten [58]. Reverse transcription using Bio-Rad iScript was performed and the resulting cDNA was used as a template for qPCR to measure relative transcript amounts. Quantitative RT-PCR data was normalized using *UBQ10* and *GAPDH* as housekeeping gene controls. Reactions were performed in a Bio-Rad CFX96 thermocycler using 96 well plates and a reaction volume of 20μL/well. SYBRGreen mastermix (Bio-Rad) was used in all reactions. Three biological and three technical replicates were used for each amplification. Primers are listed in Table S1. The MIQE guidelines were followed [59] and primer efficiencies are listed in Table S3. The thermocycling program for all RT-qPCR was a ten-minute denaturing step at 95° followed by 45 cycles of 10s at 95°, 15s at 60°, and 13s at 72°. Following amplification, melt curve analysis was done on all reactions to ensure target specificity. The melt curve program for all RT-qPCR was from 65°-95° at 0.5° increments for 5s each.

### 4.4 Repeat recombination qPCR

DNA was collected from the mature leaves of Columbia-0 and generation ten *ung* plants using the CTAB DNA extraction method [60]. qPCR was performed using primers from the flanking sequences of the intermediate repeats. Primers are listed in Table S1. Using different combinations of forward and reverse primers, either the parental or recombinant forms of the repeat can be selectively amplified (see Figure 2a). The mitochondrially-encoded *cox2* and *rrn18* genes were used as standards for analysis. Reactions were performed in a Bio-Rad CFX96 thermocycler using 96 well plates with a reaction volume of 20μL/well. SYBRGreen mastermix (Bio-Rad) was used in all reactions. Three biological and three technical replicates were used for each reaction. The thermocycling program for all repeat recombination qPCR was a ten-minute denaturing step at 95° followed by 45 cycles of 10s at 95°, 15s at 60°, and a primer specific amount of time at 72° (extension times for each primer pair can be found in Table S2). Following amplification, melt curve analysis was done on all reactions to ensure target specificity. The melt curve program for all qPCR was from 65°-95° at 0.5° increments for 5s each.

### 4.5 DNA sequencing

DNA extraction from frozen mature leaves of Columbia-0, generation 10 and *ung*, and MTP-A3G plants, and from young and mature leaves of the progeny of these plants was done by a modification of the SPRI magnetic beads method of Rowan *et al* [61, 62]. Genomic libraries for paired-end sequencing were prepared using a modification of the Nextera protocol [63], modified for smaller volumes following Baym et al [64]. Following treatment with the Nextera Tn5 transpososome 14 cycles of amplification were done. Libraries were size-selected to be between 400 and 800bp in length using SPRI beads [62]. Libraries were sequenced with 150bp paired-end reads on an Illumina HiSeq 4000 by the Vincent J. Coates Genomics Sequencing Laboratory at UC Berkeley.

Reads were aligned using BWA-MEM v0.7.12-r1039 [34]. The reference sequence used for alignment was a file containing the improved Columbia-0 mitochondrial genome (accession BK010421.1) [65] as well as the TAIR 10 *Arabidopsis thaliana* nuclear chromosomes and chloroplast genome sequences [66]. A large portion of the mitochondrial genome has been duplicated into chromosome 2 [67]. To prevent reads from mapping to both locations, this large NuMT region was deleted from chromosome 2. Using Samtools v1.3.1 [68], bam files were sorted for uniquely mapped reads for downstream analysis.

Organellar variants were called using VarDict [35]. To minimize the effects of sequencing errors and reduce false positives, SNPs called by VarDict were filtered by the stringent quality parameters of Qmean ≥ 30, MQ ≥ 30, NM ≤ 3, Pmean ≥ 8, Pstd = 1, AltFwdReads ≥ 3, and AltRevReads ≥ 3. When calling low frequency SNPs, it is difficult to remove all false positives without also removing some true positives. By treating all samples to the same sequence analysis pipeline, all samples will have a similar spectrum of false positives. By analyzing the ratios of different SNP types, rather than raw SNP numbers, we further isolate biological effects from computational noise.

DNA damage during library preparation was measured by individually analyzing the paired ends of Illumina paired end sequencing and looking for imbalances in G to T mutations between the paired ends [42]. Mapped bam files were split into separate pairs using Samtools view and analyzed with VarDict as described above. Due to the reduction in read depth by ½ while working with only one set of paired-ends, the quality filter was adjusted so that AltFwdReads ≥ 1 and AltRevReads ≥ 1. All other quality parameters remained the same. For each sample, the number of G to T mutations called in the R1 reads was divided by the number of G to T mutations called in the R2 to get a Global Imbalance Value (GIV_G_T_). A GIV_G_T_ above 1.5 is considered damaged.

Nuclear variants were called using Samtools mpileup (v. 1.3.1) and Bcftools call (v. 1.2) and filtered for SNPgap of 3, Indelgap of 10, RPB>0.1 and QUAL>15, at least 3 high quality ALT reads (DP4[2]+DP4[3] ≥3), at least one high quality ALT read per strand (DP4[2] ≥ 1 and DP4[3] ≥ 1), and a high quality ALT allele frequency ≥ 0.3. To avoid false positives, a 5 Mb region of each chromosome was used for analysis, avoiding centromeric and telomeric regions. Regions used for this analysis were: Chromosome 1 2Mb-7Mb, chromosome 3 3Mb-8Mb, chromosome 4 7Mb-12Mb, chromosome 5 3Mb-8Mb. Chromosome 2 was excluded from this analysis to avoid false positives resulting from the presence of the large NuMT that has been duplicated and repeated there.

## Supplementary Materials

The following are available online at www.mdpi.com/xxx/s1, Supplemental file 1 containing Figure S1, Table S1, Table S2 and Table S3, and Supplemental file 2 containing SNP analysis tables.

## Author Contributions

All authors have read and agree to the published version of the manuscript. Conceptualization, E.L.W. and A.C.C.; methodology, E.L.W., E.P. and A.C.C.; software, E.L.W.; validation, E.L.W. and E.P.; formal analysis, E.L.W.; investigation, E.L.W. and E.P.; resources, A.C.C.; data curation, A.C.C.; writing—original draft preparation, E.L.W. and A.C.C.; writing—review and editing, E.L.W., E.P. and A.C.C.; visualization, E.L.W.; supervision, A.C.C.; project administration, A.C.C.; funding acquisition, A.C.C.

## Acknowledgments

Conversations with Arnie Bendich about organelle DNA replication and repair in meristem and vegetative cells were interesting and illuminating. We are grateful to Emily Jezewski for finding time in her busy golf schedule to do some of the qPCR experiments. We thank Christian Elowski and the Nebraska Center for Biotechnology Core Research Facility for Microscopy for confocal fluorescent microscopy. This work used the Vincent J. Coates Genomics Sequencing Laboratory at UC Berkeley, supported by NIH S10 OD018174 Instrumentation Grant. Daniel Schachtman helped with disposal of leaf tissues from generations 2-9. The use of product and company names is necessary to accurately report the methods and results; however, the United States Department of Agriculture (USDA) neither guarantees nor warrants the standard of the products, and the use of names by the USDA implies no approval of the product to the exclusion of others that may also be suitable. The USDA is an equal opportunity provider and employer.

