## Supplemental File 1 for "Mitochondrial DNA Repair in an *Arabidopsis thaliana* Uracil N-Glycosylase Mutant"

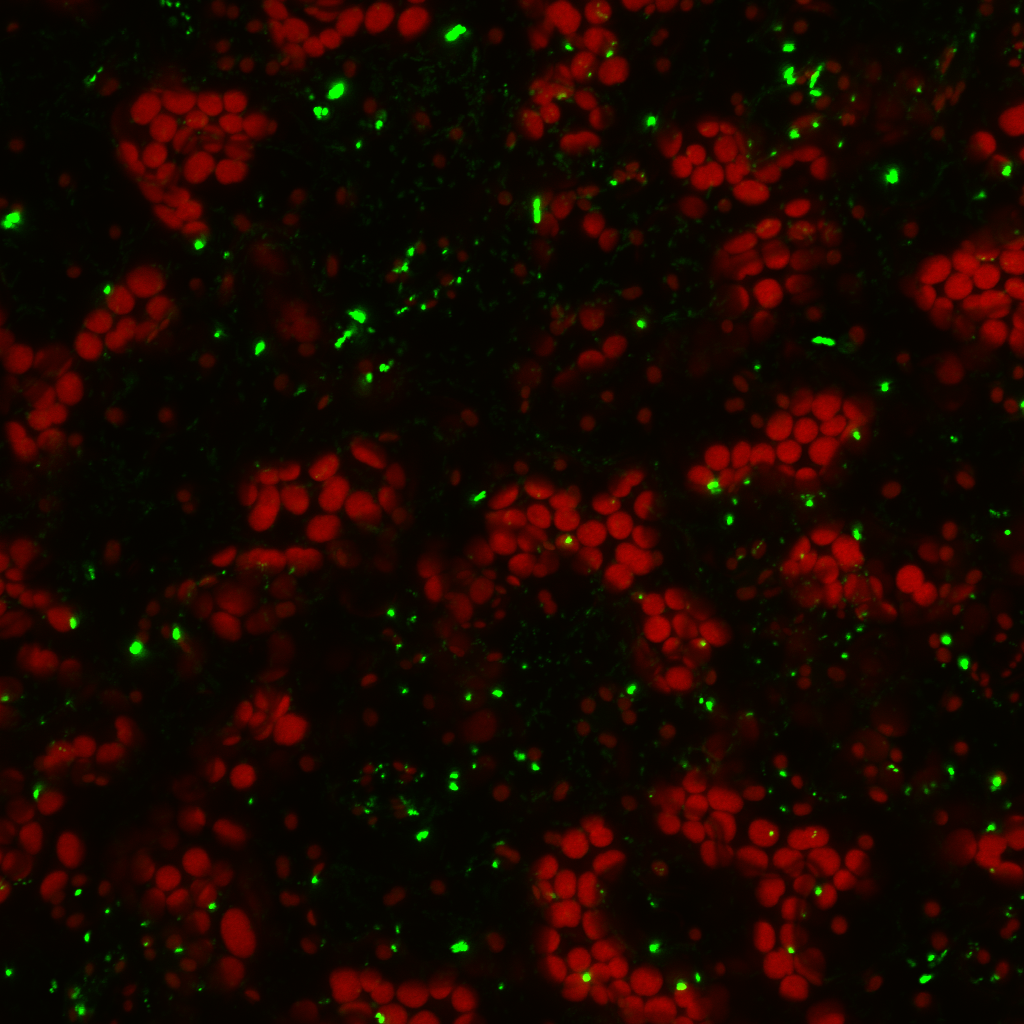

**Supplemental Figure S1: Mitochondrial targeting of a GFP labeled MTP-APOBEC3G construct.** Fluorescence microscopy of an *Arabidopsis thaliana* plant transformed with an MTP-APOBEC3G-GFP construct. Green mitochondria indicate the proper expression and targeting of the construct. Autofluorescence of chloroplasts can be seen in red.

Supplemental Table S1: Primers for RT-PCR. ^ represents exon junctions.

| Name | Size | Forward Primer | Reverse Primer |
| --- | --- | --- | --- |
| UBQ10 | 70 | TCTCAACCGTGATCAAG^ATGC | TTCCACCTCGAGGGTGATAGT |
| MSH1 | 77 | AGTTACCAAGCTCCGCACTC | CACGTC^CCTGATGCATTGTG |
| RECA2 | 74 | GCGAAGAG^ACAACGGGAAGT | GAGCAGTTCTAAACGGCGGA |

Supplemental Table S2: Primers for ROUS recombination assay

| Repeat Name | Size | Forward Primer | Reverse Primer | Extension time during qPCR |
| --- | --- | --- | --- | --- |
| Repeat L1 | 248 | GTTATGCCATTTTGGGCTTTGCTC | CGCTCGACCGAAGAAATGAGTAAC | 16s |
| Repeat L2 | 248 | CCGGTTGAAAGCTAAGCCGGAG | CCCTCACTGAACCGACTTGAATCTG | 16s |
| Repeat B1 | 532 | AAGTCCTGCTCATATACATAC | AATCCGGCTCGCTCTCGGCTACGTG | 30s |
| Repeat B2 | 532 | TACGGCTTACGAAGTTGATCG | CGACATGAAAAGACGATGTCC | 30s |
| Repeat D1 | 452 | ATAAGAGAGCAAATGATGGTATAGC | GCTGCTTTTAGGAGAGTGATCTG | 30s |
| Repeat D2 | 452 | CAAAGCAAAGGATGGTTCAAC | CGATTGTATCCGACTATTGAGTG | 30s |
| cox2 | 76 | AATAAACGTGATTGACCCAATTCT | TCCGATGAGCAGTCACTCAC | 6s |
| rrn18 | 62 | CCTTGAGCTAGGAGCCTCTTT | CATGCAAGTCGAACGTTGTT | 6s |

Supplemental Table S3: qPCR primer efficiency

| Primer Name | Calibration Curve | R^2^ | Efficiency | Limit of Detection | No Template Control Results |
| --- | --- | --- | --- | --- | --- |
| cox2 | y=-3.281x+16.06 | 0.9996 | 101.74 | 32.28 | No amplification in 45 cycles |
| rrn18 | y=-3.25x+17.295 | 0.9987 | 103.09 | 30.09 | Average Cq = 35.65. Melt curve shows nonspecific amplification |
| L11 | y=-3.681x+20.892 | 0.9979 | 86.92 | 31.5 | No amplification in 45 cycles |
| L22 | y=-3.264x+21.288 | 0.985 | 102.48 | 31.82 | No amplification in 45 cycles |
| L12 | y=-3.706x+21.916 | 0.9963 | 86.14 | 33.02 | No amplification in 45 cycles |
| L21 | y=-3.48x+24.28 | 0.9838 | 93.83 | 30.96 | No amplification in 45 cycles |
| B11 | y=-3.76x+20.928 | 0.9981 | 84.48 | 31.87 | No amplification in 45 cycles |
| B22 | y=-3.625x+20.355 | 0.9993 | 88.74 | 27.66 | No amplification in 45 cycles |
| B12 | y=-4.285x+25.995 | 0.9985 | 71.15 | 34.66 | No amplification in 45 cycles |
| B21 | y=-3.445x+23.345 | 0.9998 | 95.11 | 30.26 | No amplification in 45 cycles |
| D11 | y=-3.902x+21.65 | 0.9981 | 80.42 | 37.35 | No amplification in 45 cycles |
| D22 | y=-4.013x+21.033 | 0.9975 | 77.5 | 33.32 | No amplification in 45 cycles |
| D12 | y=-4.045x+27.638 | 0.9998 | 76.69 | 35.76 | No amplification in 45 cycles |
| D21 | y=-3.655x+26.272 | 1 | 87.76 | 33.57 | No amplification in 45 cycles |
| ubq10 | y=-3.579x+20.534 | 0.9991 | 90.29 | 34.88 | No amplification in 45 cycles |
| msh1 | y=-3.308x+33.313 | 0.9971 | 100.58 | 40.26 | Average Cq = 42.64. Melt curve shows nonspecific amplification |
| recA2 | y=-3.24x+28.263 | 0.9999 | 103.54 | 34.73 | No amplification in 45 cycles |
